# Label-Free, Physics-Constrained Learning for Multi-Exposure Speckle Imaging Parameter Estimation

**DOI:** 10.64898/2026.02.03.700085

**Authors:** Hengfa Lu, Jewel Ashbrook, Andrew K. Dunn

## Abstract

Multi-exposure speckle imaging (MESI) estimates flow-related parameters by fitting a physics-based speckle contrast model to measurements acquired over multiple exposure times. In standard pipelines, parameters are recovered via nonlinear least-squares fitting at each pixel, which is computationally expensive and can yield spatially inconsistent maps when uncertainty in the estimated speckle contrast variance 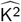 (from camera noise and finite spatial/temporal sampling used to compute speckle contrast) is amplified by independent pixel wise inversion. This work reframes MESI parameter estimation as identification of a globally shared inverse operator of the analytical forward model, exploiting the fact that a single physical mapping governs all pixels while noise drives large variance in independent pixel wise inversion. Rather than solving millions of iterative optimizations, a single parameterized inverse mapping is learned directly from a single acquired MESI dataset. Physics consistency is enforced by embedding the fixed MESI forward model as an analysis-by-synthesis layer that re-synthesizes speckle contrast curves from the predicted parameters. Training is self-supervised: the inverse mapping is optimized by minimizing a reconstruction loss between measured and re-synthesized speckle contrast curves, which constrains estimates to the set of physically admissible MESI curves without requiring ground truth parameter labels. Experiments on a numerical MESI phantom with known ground truth and on *in vivo* mouse cortex data show that the proposed method produces more stable inverse correlation time (ICT) maps (1*/τ*_c_) and improved spatial coherence relative to conventional per-pixel fitting, while substantially reducing inference time by replacing iterative optimization with a single feed-forward evaluation.

## I. Introduction

IMAGING of blood flow is essential for a wide variety of physiological measurements in the brain and other tissues [1], [2]. Laser speckle contrast imaging (LSCI) is widely used because it provides wide field, high resolution maps of speckle contrast at video rates with simple instrumentation [3]–[6]. However, single exposure LSCI is primarily qualitative: the measured speckle contrast depends not only on flow but also on static scattering, sampling statistics, and instrument-dependent factors, which limits quantitative interpretation in *in vivo* settings [7], [8].

Multi-exposure speckle imaging (MESI) addresses these limitations by acquiring speckle contrast over multiple exposure times and fitting an analytical forward model that explicitly parameterizes contributions from dynamic scattering, static scattering, and instrumentation [9]–[14]. In standard pipelines, MESI parameters are recovered by solving an independent nonlinear least-squares (NLLS) problem at each pixel. While conceptually straightforward, per-pixel NLLS has two practical drawbacks. First, iterative optimization repeated over the full field of view is computationally expensive. Second, the pixel-wise formulation is statistically inefficient: each pixel is inverted in isolation even though every pixel is governed by the same forward mapping and the same acquisition protocol. Under practical acquisition conditions, estimation error in 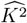, driven by camera noise and by the finite spatial/temporal sampling used to compute local contrast, propagates through independent per-pixel inversion, yielding elevated pixel-to-pixel variance and spatially inconsistent parameter maps [15]– [18].

Related efforts to accelerate or stabilize quantitative MESI analysis fall into three broad directions. First, the standard pipeline solves a nonlinear least-squares problem independently at each pixel; in principle, stability can be improved by incorporating an explicit spatial prior (for example, total-variation regularization), at the cost of converting pixel-wise fitting into a large-scale, coupled optimization problem [19]. Second, supervised learning methods have been explored to map multi-exposure speckle measurements to flow-related quantities using simulated or proxy labels, but practical deployment is constrained by the lack of pixel-wise *in vivo* ground truth and by sensitivity to mismatch between training data and experimental measurements [20]–[24]. Third, generic self-supervised denoising approaches reduce the need for clean targets in bioimaging [25]–[28], but they do not address MESI parameter inversion and do not enforce consistency with the analytical MESI forward model.

Motivated by this gap, this paper proposes a self-supervised, physics-constrained approach that learns a globally shared inverse operator of the analytical MESI forward model directly from a single acquired MESI dataset, without requiring ground-truth parameter labels. A compact, parameterized inverse mapping predicts per-pixel parameters from measured speckle contrast curves (or local MESI patches). Physical consistency is enforced by embedding the fixed, differentiable MESI forward model as an analysis-by-synthesis layer that re-synthesizes speckle contrast curves from the predicted parameters. Learning is self-supervised: the inverse mapping is optimized by minimizing a reconstruction loss between measured and re-synthesized speckle contrast curves. This objective restricts estimates to forward-model-consistent MESI curves, while the globally shared inverse operator aggregates information across the dataset to suppress estimator variance relative to independent pixel-wise inversion. At inference, parameter mapping is obtained via a single forward pass, replacing per-pixel iterative optimization.

The main contributions are as follows:

- *A global inverse formulation for MESI*. MESI parameter estimation is reformulated from millions of independent per-pixel nonlinear fits into identification of a single, globally shared inverse operator of the analytical MESI forward model, reflecting the fact that the same physics and exposure protocol govern all pixels.
- *Physics-constrained, label-free learning by analysis-by-synthesis*. A self-supervised identification strategy is in-
- troduced in which the inverse mapping is learned directly from a single acquired MESI dataset by minimizing mismatch between measured speckle contrast curves and curves re-synthesized by the fixed, differentiable MESI forward model, thereby enforcing forward-model consistency without ground-truth parameter labels.
- *Statistical efficiency and computational scaling*. By sharing a single inverse operator across the field of view
- and constraining it with the forward model, the proposed method aggregates information across the dataset to suppress noise-driven estimator variance, while reducing inference to a single forward evaluation that replaces iterative per-pixel optimization.
- *Controlled and in vivo validation*. Numerical phantom experiments with known ground truth and *in vivo* mouse
- cortex experiments demonstrate improved stability and spatial coherence of inverse correlation time (ICT) maps (1*/τ*_*c*_) relative to per-pixel nonlinear least-squares fitting, together with substantial runtime reduction.

## II. Methods and Theory

### A. MESI forward model and notation

A MESI acquisition records raw speckle intensity images at a set of exposure times 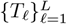. From these raw images,the squared speckle contrast is computed at each pixel using local spatial statistics [9], [10], [14]. For pixel *p*, we denote the resulting finite-sample estimates as

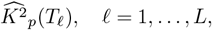

and collect them into a length-*L* vector

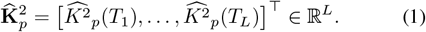

The sequence 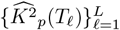 is referred to as the speckle-contrast curve at pixel *p*.

Under constant illumination, the analytical MESI forward model relates the expected squared speckle contrast to a per-pixel parameter vector

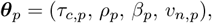

where *τ*_*c*_ is the decorrelation time (flow-related index), *ρ* ∈ [0, 1] is the dynamic scattering fraction, *β* ∈ (0, 1] is the coherence factor, and *vn* ≥0 is an additive variance offset that captures contributions not represented by the idealized model. Let *x*_*ℓ*_ = *T*_*ℓ*_*/τ*_*c,p*_ and define

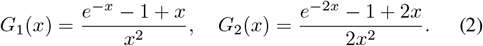

The MESI forward model can then be written as

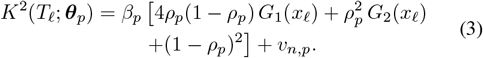

For compactness, define the forward operator

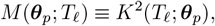

and the corresponding forward-model speckle-contrast curve vector

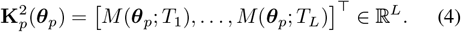

The MESI parameter estimation problem considered in this paper is to estimate ***θ***_*p*_ from 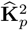 for all pixels *p*.

### B. Conventional per-pixel nonlinear least-squares fitting

Given 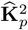 and the analytical forward operator *M* (***θ***_*p*_ ; *T*_*ℓ*_ ) in (3), the standard MESI estimator recovers parameters by solving a separate nonlinear least-squares problem at each pixel [9], [10], [14]:

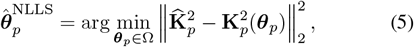

where Ω enforces physical constraints (e.g., *τ*_*c,p*_ *>* 0, *ρ*_*p*_ ∈ [0, 1], *β*_*p*_ ∈ (0, 1], and *vn*_,*p*_ ≥0). In practice, (5) is solved iteratively using local second-order methods, commonly in a trust-region or Levenberg-Marquardt framework (e.g., [29], [30]), and the procedure is repeated independently for all pixels.

This pixel-wise formulation is straightforward, but it scales poorly with image size because it requires an iterative solver at each pixel. Moreover, under practical acquisition conditions, 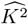 is affected by camera noise and by estimator uncertainty associated with finite spatial/temporal sampling used to compute local contrast. These perturbations are amplified by independent pixel-wise inversion, yielding parameter maps with elevated pixel-to-pixel variability. These limitations motivate an alternative formulation that replaces many independent local inversions with a single, globally shared inverse operator constrained by the MESI physics.

### C. Globally shared inverse operator with physics-constrained analysis-by-synthesis

Conventional MESI parameter estimation recovers ***θ***_*p*_ by solving a nonlinear inverse problem independently at each pixel *p*. This strategy overlooks a structural property of MESI: under a fixed acquisition protocol, the same analytical forward model governs every pixel. Consequently, shared information across the field of view is not exploited during inversion, and measurement perturbations translate into large pixel-to-pixel variability in the recovered parameters. We therefore formulate MESI parameter estimation as learning a globally shared inverse operator of the analytical MESI forward model directly from a single acquired MESI dataset.

Specifically, we introduce a parameterized inverse mapping

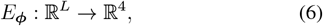

with shared parameters ***ϕ***, that maps a measured speckle-contrast curve (or a local MESI patch containing such curves) to per-pixel parameter estimates. For each pixel *p*,

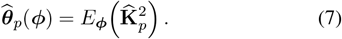

The key distinction from per-pixel NLLS is that ***ϕ*** is shared across all pixels, so the inverse mapping is learned by aggregating information across the dataset rather than estimating parameters from each pixel in isolation.

Physical consistency is enforced via an analysis-by-synthesis construction. Given 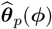, the fixed MESI forward model synthesizes the corresponding speckle-contrast curve,

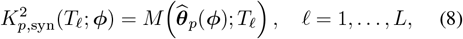

and we collect these values as

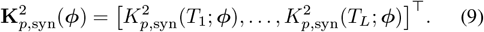

Training is self-supervised and does not require ground-truth parameter labels. Instead, ***ϕ*** is optimized by minimizing a reconstruction loss between measured and synthesized speckle-contrast curves, aggregated over pixels (or over batches of local patches),

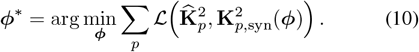

We optimize the shared inverse mapping using a reconstruction loss defined in log space. Specifically, we use a mean squared logarithmic error (MSLE) across exposure times,

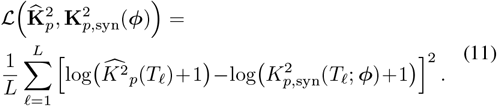

This log-space objective emphasizes relative discrepancies across exposure times and tissue types, preventing the fit from being dominated by high-contrast points while retaining sensitivity in low-*K*2 (fast-flow) regimes.

Because *M* is fixed and differentiable, optimization can reduce Eq. (10) only by adjusting 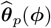 so that its forward projection under *M* matches the observed speckle-contrast curve. This couples the learned inverse mapping directly to the analytical MESI model and suppresses solutions that are inconsistent with the measurement physics.

From an estimation viewpoint, Eq. (10) replaces *N*pixels independent nonlinear optimizations with a single global learning problem for a shared inverse operator. The shared parameterization leverages redundancy across pixels and exposures, reducing variance relative to independent pixel-wise inversion under the same acquisition conditions. In practice, ***ϕ*** is optimized by stochastic gradient methods on randomly sampled local patches: each iteration evaluates *E*_***ϕ***_, synthesizes curves via *M*, computes Eq. (11), and backpropagates through *M* to update ***ϕ***. After training, inference reduces to applying 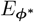 once to the full MESI stack to obtain (*τ*_*c*_, *ρ, β, vn*) maps.

### D. Parameterized inverse operator and feasibility constraints

The globally shared inverse operator in Section II-C is realized as a differentiable, parameterized mapping *E*_***ϕ***_. In principle, *E*_***ϕ***_ may be represented by different function classes, ranging from pixel-wise multilayer perceptrons to fully convolutional models operating on the multi-exposure stack. In this work we adopt a compact fully convolutional parameterization. This choice follows from the structure of MESI data: under a fixed acquisition protocol, the forward physics is identical across pixels, while the parameter fields are locally correlated (vascular continuity and smoothly varying background). A small CNN introduces an explicit locality bias and translation equivariance, improving statistical efficiency when learning from a single acquired dataset, while limiting capacity to reduce overfitting in a label-free setting.

We apply *E*_***ϕ***_ convolutionally to the *L*-channel speckle-contrast stack (or to randomly sampled local patches during training). At each pixel *p*, the network outputs intermediate variables **z***p* ∈ ℝ^4^, which are deterministically mapped to the physical parameter estimates 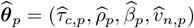. The fully convolutional construction enables efficient training from patches and, after convergence, a single forward evaluation over the full field of view.

Because the objective in Eq. (10) supervises *E*_***ϕ***_ only through curve reconstruction, feasibility must be enforced explicitly. Without constraints, optimization can enter non-physical regimes (e.g., *ρ* ∉ [0, 1] or *vn <* 0), where the forward model is not meaningful and gradient-based learning becomes unstable. We therefore impose feasibility by construction via monotone transforms that map **z***p* into a physically admissible domain:

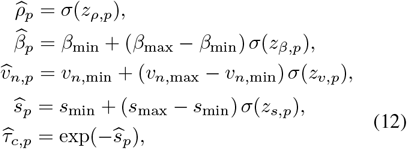

where *σ*(*·*) is the logistic sigmoid. We parameterize decorrelation through *s* = log(1*/τ*_*c*_) to accommodate the wide dynamic range of correlation times and to improve conditioning during optimization. The bounds (*β*_min_, *β*_max_), (*vn*_,min_, *vn*_,max_), and (*s*min, *s*max) are set conservatively to cover the range supported by the exposure schedule and typical operating conditions (Section II-E).

### E. Implementation details

Unless stated otherwise, the inverse operator *E*_***ϕ***_ is identified from a single measured MESI dataset and then applied to the same dataset for inference. For each dataset, we train on randomly sampled *h × w × L* patches extracted from the measured 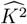 stack. The network input is globally normalized by the maximum value of 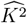 in the stack, while the reconstruction target remains in the original scale so that the analysis-by-synthesis loss in (11) is evaluated on physically meaningful *K*2 values.

#### 1) Network architecture

*E*_***ϕ***_ is implemented as a lightweight fully convolutional network with three layers: two 3 × 3 convolutions with 64 and 32 feature channels (each followed by ReLU), followed by a 1 *×* 1 convolution that outputs the four channels **z***p* in (12). Training is patch-based with *h* = *w* = 16 and mini-batch size 8; after training, the same network is evaluated convolutionally on the full *L*-channel stack in a single forward pass.

#### 2). Parameter bounds and optimization

The bounds in (12) are set conservatively to cover typical MESI operating conditions under the chosen exposure schedule. In all exper-iments we set *β* ∈ [0.05, 1], *ρ* ∈ [0.05, 1], and 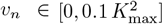, where 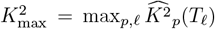 is computed from the dataset. The log-decorrelation rate is bounded by 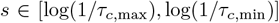 with *τ*_*c*,min_ = 10^−5^ s and 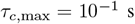

We optimize (10) using Adam (*β*_1_ = 0.9, *β*_2_ = 0.999) [31] with an initial learning rate of 10^−3^. Training is performed for 10 epochs in PyTorch on a single NVIDIA A100 GPU.

## III. Experimental Results

We evaluate the proposed globally learned, physics-constrained inverse operator on (i) a numerical MESI phantom and (ii) *in vivo* mouse cortex data. The numerical phantom provides a controlled setting with known parameter fields (phantom truth) and access to both noise-free and noise-corrupted 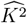 stacks, enabling direct assessment of spatial fidelity in the recovered inverse correlation time (ICT) maps. The *in vivo* experiments probe robustness under practical acquisition conditions and assess whether the proposed estimator yields anatomically coherent ICT maps relative to conventional per-pixel nonlinear least-squares (NLLS) fitting. Unless otherwise stated, all methods operate on the same 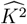 stacks, computed with identical local contrast-estimation settings, and use the same 15-exposure schedule.

### A. Experimental Setups

#### 1) Numerical MESI phantom

We constructed a numerical MESI phantom to enable controlled evaluation in the speckle-contrast and ICT domains. The phantom is defined by spatially varying parameter fields ***θ***(*p*) = (*τ*_*c,p*_, *ρ*_*p*_, *β*_*p*_, *vn*_,*p*_), which are used as ground-truth maps for synthesis and evaluation. To obtain physiologically realistic spatial structure, these fields were derived from high-SNR *in vivo* reference estimates (obtained by fitting heavily averaged speckle-contrast data) and then treated as fixed for phantom generation.

Given the exposure schedule 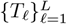 with *L* = 15, we generated a noise-free squared speckle-contrast stack by evaluating the analytical MESI forward model (3) at each pixel and exposure time. To emulate realistic acquisition, we synthesized noise-corrupted measurements by generating repeated noisy raw speckle intensity realizations with a camera-like noise model (Poisson shot noise and additive Gaussian read noise), computing 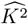 for each realization using the same local spatial estimator as in experiments (window size *w × w*), and then averaging the resulting 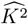 images across *N*rep realizations at each exposure. This procedure mirrors practical MESI preprocessing (averaging in the speckle-contrast domain rather than averaging raw intensities) and yields matched triplets for evaluation: (i) phantom-truth parameter maps, (ii) a noise-free 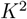 stack, and (iii) a noise-corrupted 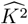 stack. Unless otherwise stated, we use *w* = 7, mean intensity *µ*_*I*_ = 1600 electrons/pixel, read-noise standard deviation *σ*_*r*_ = 0.5 electrons, and *N*rep = 150.

#### 2) In vivo mouse cortex imaging

*In vivo* MESI data were acquired with a standard epi-illumination configuration. A 785 nm laser diode (L785P090, Thorlabs) provided coherent illumination, expanded to uniformly illuminate the exposed cortical surface. Back-scattered light was collected by a camera lens (AF NIKKOR 50 mm f/1.8D, Nikon) and imaged onto a monochrome CMOS camera (acA1920-155um, Basler). For each dataset, raw speckle intensity frames were acquired at *L* = 15 exposure times using the same schedule as the numerical phantom (log-spaced from 50 *µ*s to 80 ms). At each exposure, squared speckle contrast 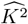 was computed at each pixel using local spatial statistics [9], [10], [14], and 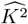 images were averaged across repeated realizations to form the input stack used for parameter estimation.

All animal procedures were approved by the Institutional Animal Care and Use Committee (IACUC) of the University of Texas at Austin. Experiments were performed on adult male CD-1 mice (25–30 g, Charles River Laboratories). Mice were anesthetized with 2–3% isoflurane in a 70% N2/O2 mixture, and body temperature was maintained at 37.5^◦^C with a feedback-controlled heating system (FHC, Bowdoin, ME).

### B. Numerical Phantom Validation

We first verify that the noise-corrupted phantom measurements remain within a realistic perturbation regime for MESI fitting. Figure 2 shows representative *K*2-versus-exposure curves at four locations, comparing the noise-free forward-model values with the corresponding noise-corrupted measurements. Across the exposure range, the corrupted curves track the noise-free curves closely, indicating that the phantom captures realistic finite-sample variability without qualitatively altering the exposure dependence dictated by the forward model.

**Fig. 1.**
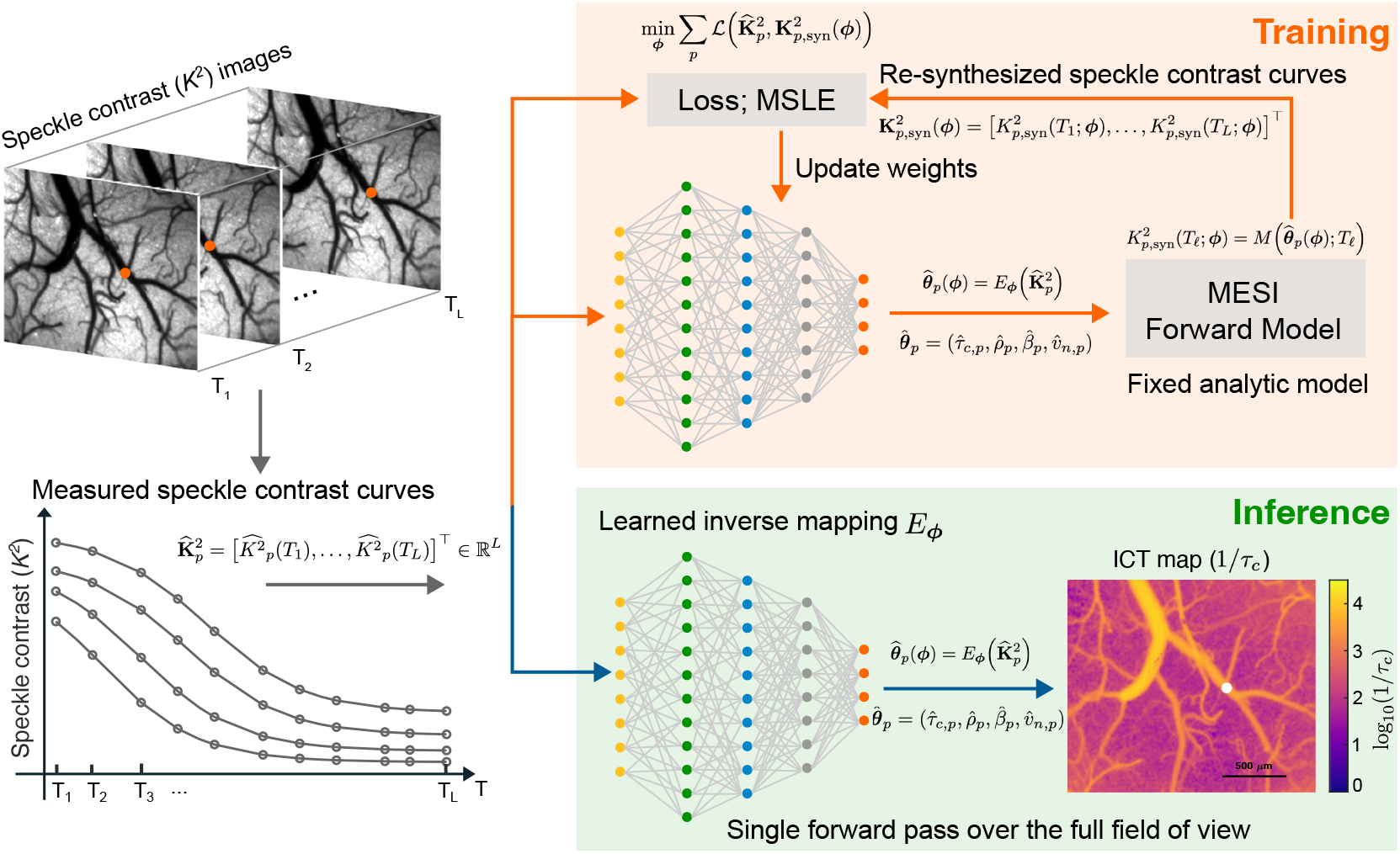
Physics-constrained analysis-by-synthesis learning of a globally shared inverse operator for MESI. During training, a parameterized inverse mapping ***E***_***ϕ***_ predicts per-pixel parameters from measured speckle contrast curves (or local MESI patches). The fixed analytical MESI forward model ***M*** then re-synthesizes speckle contrast curves from the predicted parameters, and ***ϕ*** is optimized by minimizing a curve-reconstruction loss aggregated over pixels (or patches). At inference, parameter mapping is obtained by a single forward pass through 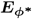, replacing per-pixel iterative optimization.

**Fig. 2.**
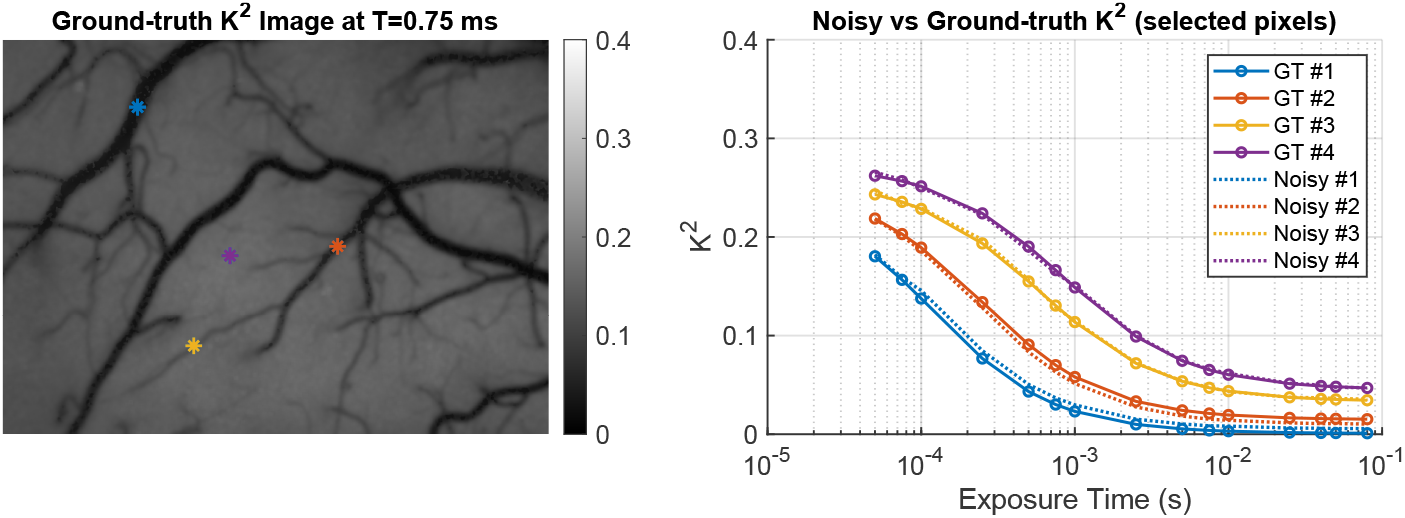
Representative speckle-contrast curves in the numerical phantom. Squared speckle contrast ***K*2** versus exposure time ***T*** at four phantom locations. Solid traces: noise-free forward-model values from (3) evaluated using the phantom-truth parameters. Dotted traces: noise-corrupted measurements formed by computing 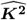 per realization from simulated raw speckle frames and averaging 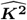 across realizations at each exposure. The corrupted curves remain close to the noise-free curves across exposures, consistent with the practical variance-reduced regime used for MESI fitting.

We next evaluate spatial fidelity in the recovered flow index. Figure 3 compares ICT maps (1*/τ*_*c*_), displayed as log_10_(1*/τ*_*c*_) for visualization, estimated by per-pixel NLLS and by the proposed estimator against the phantom truth. Per-pixel NLLS exhibits pronounced pixel-scale variability that disrupts vessel interiors and introduces localized discontinuities along vessel boundaries. In contrast, the proposed estimator yields spatially more coherent ICT maps while preserving vessel edges and small branches.

**Fig. 3.**
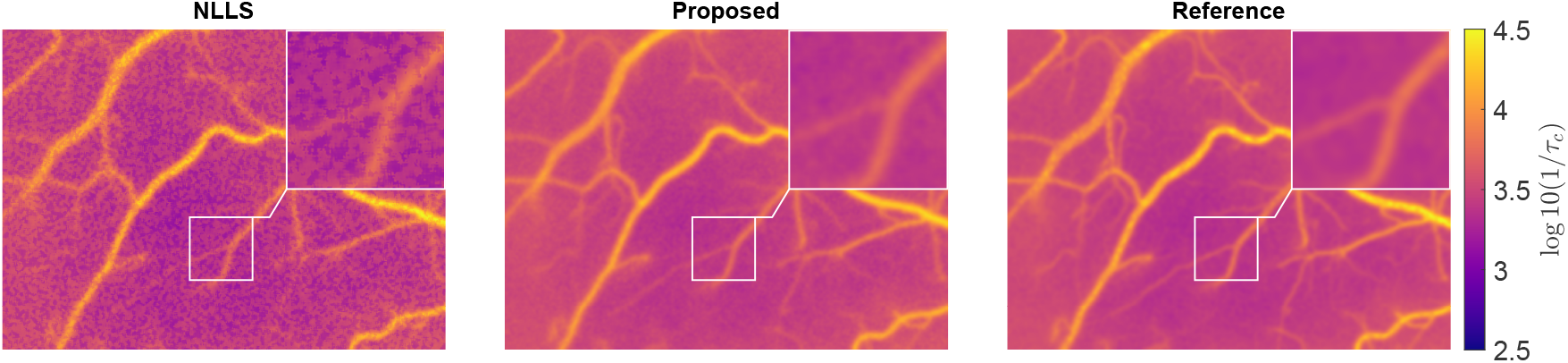
Numerical phantom ICT maps. **log_10_(1*/τ***_***c***_) reconstructed by per-pixel NLLS (left) and by the proposed estimator (middle), with phantom truth (right). Insets show the same fixed zoomed ROI (**2.5 *×***). Relative to NLLS, the proposed estimator reduces pixel-scale fluctuations while preserving vessel boundaries and small branches.

To probe local structure more directly, Fig. 4 reports a line profile crossing multiple vessels. Relative to NLLS, the proposed estimate follows the phantom-truth profile more closely along both vessel and background segments and exhibits markedly fewer high-frequency oscillations and localized over/under-shoots at vessel crossings. Together, these results support the central premise of the method: learning a globally shared inverse operator under a physics-constrained objective suppresses noise-driven variance amplification that arises when each pixel is inverted independently.

**Fig. 4.**
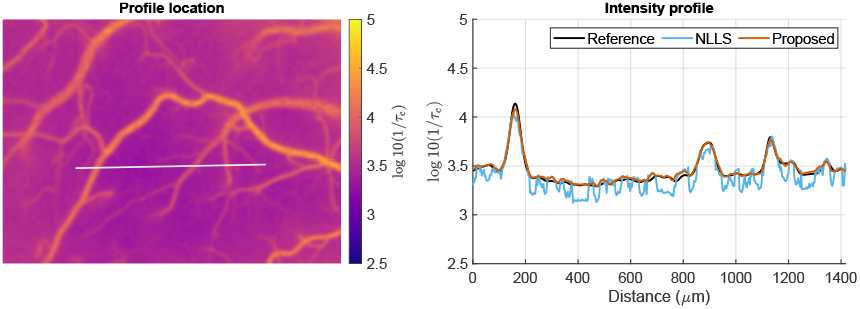
Numerical phantom line-profile analysis in the ICT domain. Left: profile location. Right: **log_10_(1*/τ***_***c***_) along the marked line for phantom truth (black), per-pixel NLLS (blue), and the proposed estimator (orange). The proposed profile more closely follows phantom truth with substantially reduced high-frequency oscillations, while preserving vessel-crossing peak locations and relative contrast.

### C. In Vivo Evaluation on Mouse Cortex

We next evaluated the proposed estimator on *in vivo* mouse cortex MESI data acquired with the system in Section III-A. Figure 5 compares ICT maps, displayed as log_10_(1*/τ*_*c*_), recovered by the conventional per-pixel NLLS baseline and by the proposed globally learned inverse operator from the same measured 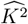 stack. The NLLS map shows elevated pixel-to-pixel variability within vessels, discontinuities along vessel boundaries, and spatially incoherent fluctuations in parenchyma-patterns consistent with independent pixel-wise inversion amplifying measurement and estimator uncertainty. In contrast, the proposed estimator yields an ICT map with improved within-region consistency and vessel continuity: vessel interiors exhibit reduced local variability, boundaries are less fragmented, and parenchymal regions contain fewer isolated outliers while preserving microvascular structure.

**Fig. 5.**
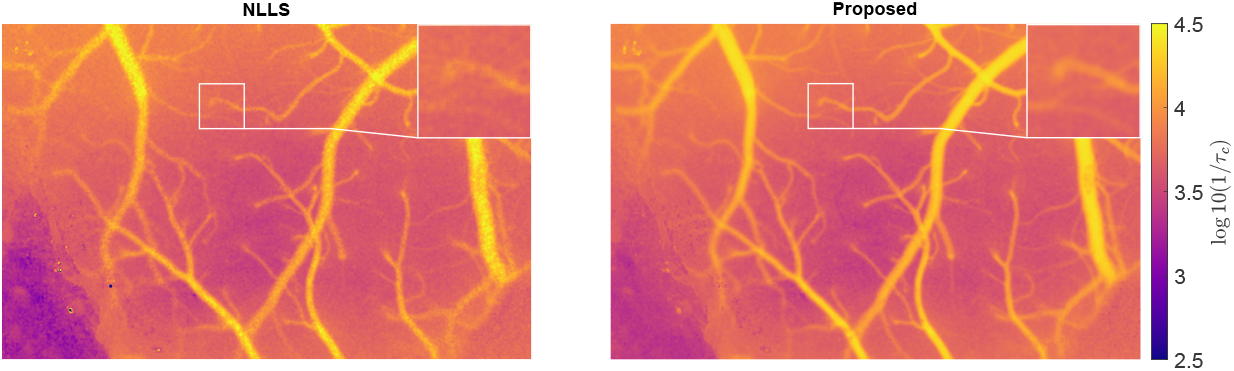
*In vivo* mouse cortex ICT maps. **log_10_(1*/τ***_***c***_) estimated by per-pixel NLLS fitting (left) and by the proposed estimator (right) from the same measured 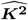 stack. Relative to NLLS, the proposed map exhibits reduced local variability within vessels, fewer boundary discontinuities, and improved spatial coherence in parenchyma while preserving vascular morphology. The overlaid line marks the profile location used in Fig. 6.

To examine spatial behavior more directly, Fig. 6 reports a line profile traversing multiple surface vessels and intervening parenchyma. Along this transect, the NLLS estimate contains high-frequency oscillations and localized over/under-shoots, particularly near vessel edges, which perturb peak amplitudes and distort vessel cross-sections. The proposed estimate exhibits a smoother profile with more stable vessel-crossing peaks and a slowly varying parenchymal baseline, consistent with reduced estimator variance while retaining the expected spatial organization of cortical flow.

**Fig. 6.**
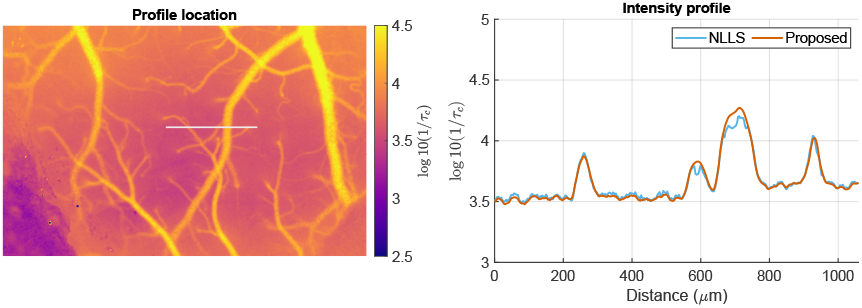
*In vivo* line-profile analysis. Left: profile location overlaid on the proposed ICT map. Right: **log_10_(1*/τ***_***c***_) along the marked line for per-pixel NLLS (blue) and the proposed estimator (orange). The proposed profile suppresses high-frequency fluctuations and yields more stable vessel-crossing peaks, indicating reduced noise-driven variability in MESI parameter estimation

Taken together, the *in vivo* results support the central claim of this work: identifying a globally shared inverse operator under a physics-constrained analysis-by-synthesis objective reduces noise-driven spatial inconsistency relative to per-pixel NLLS fitting while preserving anatomically plausible vascular structure in the recovered ICT maps.

## IV. Conclusion and Discussion

This work introduced a different formulation of quantitative MESI parameter estimation: rather than solving a separate nonlinear inverse problem at every pixel, we identify a single, globally shared inverse operator of the analytical MESI forward model directly from one acquired dataset. The inverse mapping is learned under a physics-constrained analysis-by-synthesis objective in which predicted parameters are required to reproduce the measured speckle-contrast curves through the fixed, differentiable MESI model. After training, parameter inference reduces to a single feed-forward evaluation, replacing per-pixel iterative solvers.

From an estimation perspective, the benefit follows from exploiting structure that conventional pipelines discard. Pixel-wise NLLS fitting treats each speckle-contrast curve in isolation, so perturbations in the finite-sample contrast estimates are amplified by repeated local inversions, yielding spatially inconsistent parameter maps. In contrast, a globally shared inverse operator pools information across the field of view under a common acquisition protocol, while the embedded forward model constrains the solution space to curves that are consistent with MESI physics. This coupling suppresses nonphysical degrees of freedom and reduces noise-driven variance relative to independent pixel-wise inversion.

Experiments on a controlled numerical phantom and on *in vivo* mouse cortex data support this premise. In both settings, the proposed estimator produced inverse correlation time maps with improved spatial coherence and vessel continuity relative to conventional per-pixel NLLS fitting, while preserving sharp vascular boundaries and fine structure. The approach also enables fast inference, since parameter maps are obtained by a single forward pass rather than by iterative optimization repeated over pixels.

The scope of the method is tied to the assumed measurement model and acquisition protocol. Because learning is constrained by the analytical MESI forward model, modeling mismatch (e.g., deviations from the assumed dynamics or unmodeled experimental effects) will translate into structured residuals that cannot be eliminated by the inverse mapping alone. In addition, the inverse operator is identified from a single dataset under a fixed exposure schedule; this design choice is deliberate for label-free learning, but it motivates future work on protocol-conditional or multi-dataset training to improve transfer across systems and operating conditions.

Overall, the proposed globally learned, physics-constrained inversion provides a practical alternative to conventional MESI fitting: it preserves the interpretability and consistency of analytical modeling while mitigating the variance amplification inherent to per-pixel inversion, and it replaces computationally expensive pixel-wise solvers with efficient feed-forward inference. Beyond MESI, the same principle, identifying a shared inverse operator under an analysis-by-synthesis constraint, offers a general route for stabilizing quantitative inference in imaging problems where a common forward model governs many local measurements but conventional pipelines invert them independently.

## Acknowledgment

The authors are grateful for funding from the UT Austin Portugal Program and support from the Texas Advanced Computing Center (TACC) at The University of Texas at Austin for providing computational resources that contributed to the research results reported within this paper.

## References

[1] C. Iadecola, “Neurovascular regulation in the normal brain and in Alzheimer’s disease,” Nat. Rev. Neurosci., vol. 5, no. 5, pp. 347–360, 2004.

[2] D. Attwell, A. M. Buchan, S. Charpak, M. Lauritzen, B. A. Macvicar, and E. A. Newman, “Glial and neuronal control of brain blood flow,” Nature, vol. 468, no. 7321, pp. 232–243, 2010.

[3] J. D. Briers and S. Webster, “Laser speckle contrast analysis (LASCA): a nonscanning, full-field technique for monitoring capillary blood flow,” J. Biomed. Opt., vol. 1, no. 2, pp. 174–179, 1996.

[4] A. K. Dunn, H. Bolay, M. A. Moskowitz, and D. A. Boas, “Dynamic imaging of cerebral blood flow using laser speckle,” J. Cereb. Blood Flow Metab., vol. 21, no. 3, pp. 195–201, 2001.

[5] D. A. Boas and A. K. Dunn, “Laser speckle contrast imaging in biomedical optics,” J. Biomed. Opt., vol. 15, no. 1, pp. 011 109–011 109, 2010.

[6] A. K. Dunn, “Laser speckle contrast imaging of cerebral blood flow,” Ann. Biomed. Eng., vol. 40, pp. 367–377, 2012.

[7] J. D. Briers, “Laser speckle contrast imaging for measuring blood flow,” Opt. Appl., vol. 37, 2007.

[8] D. Briers, D. D. Duncan, E. Hirst, S. J. Kirkpatrick, M. Larsson, W. Steenbergen, T. Stromberg, and O. B. Thompson, “Laser speckle contrast imaging: theoretical and practical limitations,” J. Biomed. Opt., vol. 18, no. 6, pp. 066 018–066 018, 2013.

[9] A. B. Parthasarathy, W. J. Tom, A. Gopal, X. Zhang, and A. K. Dunn, “Robust flow measurement with multi-exposure speckle imaging,” Opt. Express, vol. 16, no. 3, pp. 1975–1989, 2008.

[10] S. M. S. Kazmi, A. B. Parthasarthy, N. E. Song, T. A. Jones, and A. K. Dunn, “Chronic imaging of cortical blood flow using multi-exposure speckle imaging,” J. Cereb. Blood Flow Metab., vol. 33, no. 6, pp. 798– 808, 2013.

[11] S. S. Kazmi, S. Balial, and A. K. Dunn, “Optimization of camera exposure durations for multi-exposure speckle imaging of the microcirculation,” Biomed. Opt. Express, vol. 5, no. 7, pp. 2157–2171, 2014.

[12] S. M. S. Kazmi, R. K. Wu, and A. K. Dunn, “Evaluating multi-exposure speckle imaging estimates of absolute autocorrelation times,” Opt. Lett., vol. 40, no. 15, pp. 3643–3646, Aug 2015.

[13] C. J. Schrandt, S. S. Kazmi, T. A. Jones, and A. K. Dunn, “Chronic monitoring of vascular progression after ischemic stroke using multi-exposure speckle imaging and two-photon fluorescence microscopy,” J. Cereb. Blood Flow Metab., vol. 35, no. 6, pp. 933–942, 2015.

[14] L. M. Richards, S. S. Kazmi, K. E. Olin, J. S. Waldron, D. J. FoxJr, and A. K. Dunn, “Intraoperative multi-exposure speckle imaging of cerebral blood flow,” J. Cereb. Blood Flow Metab., vol. 37, no. 9, pp. 3097–3109, 2017.

[15] M. A. Davis, L. Gagnon, D. A. Boas, and A. K. Dunn, “Sensitivity of laser speckle contrast imaging to flow perturbations in the cortex,” Biomed. Opt. Express, vol. 7, no. 3, pp. 759–775, 2016.

[16] C. Liu, K. Kiliç, S. E. Erdener, D. A. Boas, and D. D. Postnov, “Choosing a model for laser speckle contrast imaging,” Biomed. Opt. Express., vol. 12, no. 6, pp. 3571–3583, 2021.

[17] S. Zheng and J. Mertz, “Correcting sampling bias in speckle contrast imaging,” Opt. Lett., vol. 47, no. 24, pp. 6333–6336, 2022.

[18] S. Zheng, I. Davison, A. Garrett, X. Lin, N. Chitkushev, D. Roblyer, and J. Mertz, “Robust speckle contrast imaging based on spatial covariance,” Optica, vol. 11, no. 12, pp. 1733–1741, 2024.

[19] L. I. Rudin, S. Osher, and E. Fatemi, “Nonlinear total variation based noise removal algorithms,” Phys. D: Nonlinear Phenom., vol. 60, no. 1, pp. 259–268, 1992.

[20] M. Hultman, M. Larsson, T. Strömberg, and I. Fredriksson, “Real-time video-rate perfusion imaging using multi-exposure laser speckle contrast imaging and machine learning,” J. Biomed. Opt., vol. 25, no. 11, pp. 116 007–116 007, 2020.

[21] X. Hao, S. Wu, L. Lin, Y. Chen, S. P. Morgan, and S. Sun, “A quantitative laser speckle-based velocity prediction approach using machine learning,” Opt. Laser Eng., vol. 166, p. 107587, 2023.

[22] C.-Y. Yu, M. Chammas, H. Gurden, H.-H. Lin, and F. Pain, “Design and validation of a convolutional neural network for fast, model-free blood flow imaging with multiple exposure speckle imaging,” Biomed. Opt. Express., vol. 14, no. 9, pp. 4439–4454, 2023.

[23] H.-S. Park and Y.-C. Ahn, “Method for measuring blood flow and depth of blood vessel based on laser speckle contrast imaging using 3d convolutional neural network: A preliminary study,” Opt. Laser Technol., vol. 179, p. 111367, 2024.

[24] K. J. Shang, Y. Yuan, H. L. Liu, R. B. Wang, W. N. Gao, Y. Bi, and Y. Yu, “Estimation of absolute wide-range blood flow by deep learning-based laser speckle contrast imaging,” Opt. Laser Eng., vol. 193, p. 109056, 2025.

[25] J. Lehtinen, J. Munkberg, J. Hasselgren, S. Laine, T. Karras, M. Aittala, and T. Aila, “Noise2Noise: Learning image restoration without clean data,” in Proceedings of the 35th International Conference on Machine Learning, ser. Proceedings of Machine Learning Research, J. Dy and A. Krause, Eds., vol. 80. PMLR, 10–15 Jul 2018, pp. 2965–2974.

[26] A. Krull, T.-O. Buchholz, and F. Jug, “Noise2void-learning denoising from single noisy images,” in Proc. IEEE Comput. Soc. Conf. Comput. Vis. Pattern Recognit., 2019, pp. 2129–2137.

[27] J. Batson and L. Royer, “Noise2self: Blind denoising by self-supervision,” in International conference on machine learning (ICML). PMLR, 2019, pp. 524–533.

[28] R. F. Laine, G. Jacquemet, and A. Krull, “Imaging in focus: An introduction to denoising bioimages in the era of deep learning,” Int. J. Biochem. Cell Biol., vol. 140, p. 106077, 2021.

[29] K. Levenberg, “A method for the solution of certain non-linear problems in least squares,” Quarterly of applied mathematics, vol. 2, no. 2, pp. 164–168, 1944.

[30] D. W. Marquardt, “An algorithm for least-squares estimation of non-linear parameters,” Journal of the Society for Industrial and Applied Mathematics, vol. 11, no. 2, pp. 431–441, 1963.

[31] D. P. Kingma, J. Ba et al., “Adam: A method for stochastic optimization,” in ICLR, 2015.

